# Human extracellular sulfatases use a dual mechanism for regulation of growth factor interactions with heparan sulfate proteoglycans

**DOI:** 10.1101/2023.11.22.568358

**Authors:** Bryce M. Timm, Julianna L. Follmar, Ryan N. Porell, Kimberly Glass, Bryan E. Thacker, Charles A. Glass, Kamil Godula

## Abstract

Membrane-associated heparan sulfate (HS) proteoglycans (PGs) contribute to the regulation of extracellular cellular signaling cues, such as growth factors (GFs) and chemokines, essential for normal organismal functions and implicated in various pathophysiologies. PGs accomplish this by presenting high affinity binding sites for GFs and their receptors through highly sulfated regions of their HS polysaccharide chains. The composition of HS, and thus GF-binding specificity, are determined during biosynthetic assembly prior to installation at the cell surface. Two extracellular 6-*O*-endosulfatase enzymes (Sulf-1 and Sulf-2) can uniquely further edit mature HS and alter its interactions with GFs by removing specific sulfation motifs from their recognition sequence on HS. Despite being implicated as signaling regulators during development and in disease, the Sulfs have resisted structural characterization, and their substrate specificity and effects on GF interactions with HS are still poorly defined. Using a panel of PG-mimetics comprising compositionally-defined bioengineered recombinant HS (rHS) substrates in combination with GF binding and enzyme activity assays, we have discovered that Sulfs control GF-HS interactions through a combination of catalytic processing and competitive blocking of high affinity GF-binding sites, providing a new conceptual framework for understanding the functional impact of these enzymes in biological context. Although the contributions from each mechanism are both Sulf- and GF-dependent, the PG-mimetic platform allows for rapid analysis of these complex relationships.

**Significance Statement:** Cells rely on extracellular signals such as growth factors (GFs) to mediate critical biological functions. Membrane-associated proteins bearing negatively charged heparan sulfate (HS) sugar chains engage with GFs and present them to their receptors, which regulates their activity. Two extracellular sulfatase (Sulf) enzymes can edit HS and alter GF interactions and activity, although the precise mechanisms remain unclear. By using chemically defined HS-mimetics as probes, we have discovered that Sulfs can modulate HS by means of catalytic alterations and competitive blocking of GF-binding sites. These unique dual activities distinguish Sulfs from other enzymes and provide clues to their roles in development and disease.

## Introduction

The ability of cells to sense external biochemical cues, such as growth factors (GFs) and other morphogens, is essential for normal organismal development and function.^1^ Dysregulation of GF signaling can lead to aberrant processes and emergence of disease phenotypes, including tumor formation and metastasis.^2^ The activation of GF receptors (GFRs) occurs at the cell surface in the context of the glycocalyx, a complex ensemble of glycolipids and glycoproteins that influence this process and downstream signaling.^3^ In animal cells, heparan sulfate (HS) proteoglycans (PGs) are critical glycocalyx components, regulating a myriad of signaling pathways.^4^ When shed from cells, PGs can also sequester GFs into the extracellular matrix (ECM) away from GFRs.^5^ HSPG binding of GFs and GFRs occurs through protein-specific interactions driven by the structure and sulfation of HS glycan chains.^6,7^

The structure of HS glycans and, thus GF-binding specificity, is defined during their biosynthesis in the endoplasmic reticulum and Golgi compartments. ^8^ Once presented on the cell surface, HSPGs are uniquely further modified by two extracellular 6-*O*-endosulfatase enzymes, Sulf-1 and Sulf-2, which catalyze subtle but consequential changes within GF-binding regions of the glycan chains by removing glucosamine 6-*O*-sulfates (**Fig. 1*A***).^9^ Despite being closely related isoforms, systems-level biological studies indicate specific roles for each Sulf.^10^ In cancer specifically, ^11^ over-expression of Sulf-1 can be anti-oncogenic,^12,13^ while overexpression of Sulf-2 is generally pro-oncogenic,^14^ leading to increased tumorigenesis.^15,16^ Thus, the Sulfs may present novel biomarkers for early cancer diagnosis or be the target for therapeutic intervention;^17,18^ however, how they exert their specific functions is still unclear.

**Figure 1.**
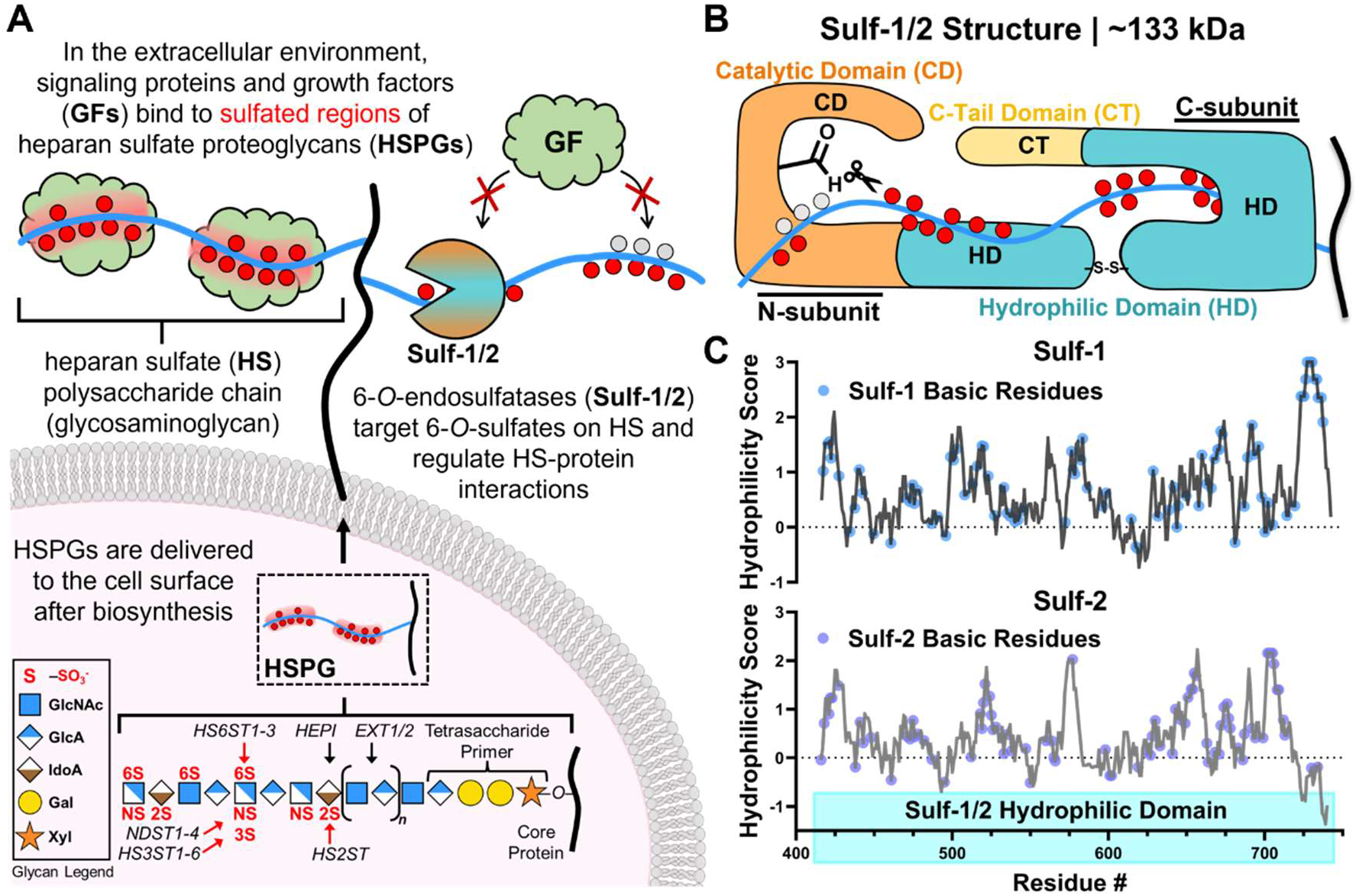
Extracellular 6-*O*-endosulfatases (Sulf-1 and Sulf-2) regulate growth factor (GF) interactions with cell-surface heparan sulfate proteoglycans (HSPGs). *(A)* In the extracellular environment, HSPGs bind GFs and other signaling proteins to regulate cellular signaling pathways. HSPGs consist of protein-linked polysaccharide chains that are modified during biosynthesis by many enzymes, resulting in interspersed regions of sulfation and structural heterogeneity. Sulfs 1 and 2 influence GF function by processive desulfation of HS through removal of specific 6-*O*-sulfation motifs. *(B)* Sulfs 1 and 2 are uniquely fashioned to bind to HS via interaction with their hydrophilic domain (HD) and catalyze 6-*O*-desulfation using a reactive formylglycine (FGly) catalytic residue in coordination with residues in the catalytic domain (CD), HD and C-tail domain (CT). (*C*) Comparison of the computed hydrophilicity of Sulf-1 and Sulf-2 hydrophilic domains and the distributions of basic residues within them.

Sulf-1 and Sulf-2 are distinct from other human sulfatases in both their localization to the ECM and in their structural complexity. While sharing a common catalytic domain (CD) with lysosomal sulfatases that contains the reactive formylglycine (FGly) residue,^19^ Sulf-1 and Sulf-2 feature an additional hydrophilic domain (HD) for the binding and processive desulfation of internal HS regions (**Fig. 1*B***).^20, 21^ A C-terminal (CT) domain completes the protein sequence and is required for enzymatic activity.^22^ The greatest sequence divergence between the two isoforms is seen in their HD region,^9^ suggesting unique HS-binding and activity profiles (**Fig. 1*C* and *SI Appendix,* Fig. S1**). However, the disordered state of the HD has impeded a full structural characterization of native Sulfs and their interactions with substrates at the molecular level.

The HS substrates for Sulfs are generated in the secretory pathway through a non-template process driven by the sequential action of glycosylation enzymes (**Fig. 1*A***). ^23^ HS chain biosynthesis initiates with the assembly of a tetrasaccharide primer on serine or threonine residues in the core protein, which is then elongated by alternating copolymerization of *N*-acetylglucosamine (GlcNAc) and glucuronic acid (GlcA) residues.^24^ The growing chains are modified by *N*- deacetylase-*N-*sulfotransferase (NDST) enzyme complexes, which modify GlcNAc through substitution with *N*-sulfate, and initiate the formation of sulfated segments. These regions are then elaborated by a C-5 epimerase (HEPI), converting GlcA residues to iduronic acids (IdoAs), and by various 2-, 6- and 3-*O*-sulfotransferases, which introduce additional sulfation to define the fine structure of HS and its protein binding function.^25,26^

The partial structural characterization of the Sulfs and complexity of their substrates have limited our understanding of their mechanism of action and biological functions. Prior studies established that the Sulfs preferentially act on trisulfated-disaccharides in sulfated HS domains,^27,28^ and that Sulf-1 processed soluble heparin oligosaccharides more efficiently than Sulf-2.^21^ However, using heparin, a highly sulfated HS analogue, as the substrate may be limited in predicting Sulf preferences for native HS structures. This limitation may be addressed by using soluble chemically defined synthetic HS oligosaccharides as substrates in *in vitro* enzymatic assays.^29^ Remarkably, a genetic mutant of A375 melanoma cells (A375 KDM2B^C5^), wherein repression of the histone demethylase KDM2B resulted in an elevated Sulf-1 expression, showed profoundly impacted GF-binding profile without a significant decrease in 6-*O*-sulfation.^30^ This suggests that Sulf-1 may regulate GF interactions and cellular activity through remarkably specific edits to HS. Such subtle changes in HS structure are, however, difficult to pinpoint using existing glycan analysis techniques and may be obscured in *in vitro* enzymatic assays.

Herein, we report our investigation into the complex relationship between HS structure, GF-binding, and the mechanisms of regulation by Sulfs. We evaluated the effects of Sulf-1 and Sulf-2 on the binding of various GFs against a panel of PG-mimetic bioconjugates generated by grafting recombinant HS (rHS) polysaccharides with defined composition and sulfation characteristics to a bovine serum albumin (BSA) protein carrier. Using a hybrid binding/activity assay, we discovered that the Sulfs utilize a dual mechanism for controlling GF interactions with HS by combining the catalytic remodeling of the sulfation domains with competitive blocking of GF-binding sites. We investigated the relative effects of these two mechanisms on the binding of fibroblast growth factor 1 and 2 (FGF1, FGF2), vascular endothelial growth factor (VEGF), and bone morphogenic protein 4 (BMP4), in relation to the sulfation status of the rHS substrates for each Sulf. We also compared the relative substrate specificity and differential activity of Sulf-1 and Sulf-2 in the context of FGF1 interactions with HS. We found that Sulf-1 exerted a stronger impact on the binding of this GF via both mechanisms, while Sulf-2 limited FGF1 binding to a lesser extent, mainly through catalytic substrate processing. Collectively, our findings provide new insights into the substrate specificity and activity of Sulfs and establish a framework for understanding their contributions to the regulation of GF signaling.

## Results and Discussion

### Generation of PG-mimetic ELISA platform for profiling Sulf activity

Since 2002, glycan array platforms have revolutionized glycomics by enabling high-throughput analyses of glycan-protein interactions.^31, 32^ Despite advances in generating glycan arrays with increasingly complex synthetic HS oligosaccharides,^33^ the size of accessible glycan chains and their accessibility to enzymatic processing after immobilization have limited the applicability of these tools for the analysis of substrate-specificity of Sulfs and their influence on HS-GF interactions. To evaluate the effects of Sulf activity on GF and chemokine binding to HS, Rosen and co-workers introduced the use of heparin and porcine intestinal mucosal HS conjugated to bovine serum albumin (BSA) as model Sulf substrates in enzyme-linked immunosorbent assays (ELISAs). ^34,35^

We recently published an improved bioconjugation strategy for generating HS-BSA PG-mimetics^36^ using various rHS polysaccharides with well-defined compositions sourced from cell lines with genetically-engineered HS biosynthesis.^37^ These BSA-linked conjugates provided accessible reagents for profiling the ligand-binding specificity of HS-binding proteins and the activity of HS-processing enzymes.^36,38^ We envision that these conjugates could be used as substrates in enzymatic and binding assays to systematically characterize the binding, catalytic activity, and isoform specificity of Sulfs in the context of their regulation of GF-HS interactions (**Fig. 2**).

**Figure 2.**
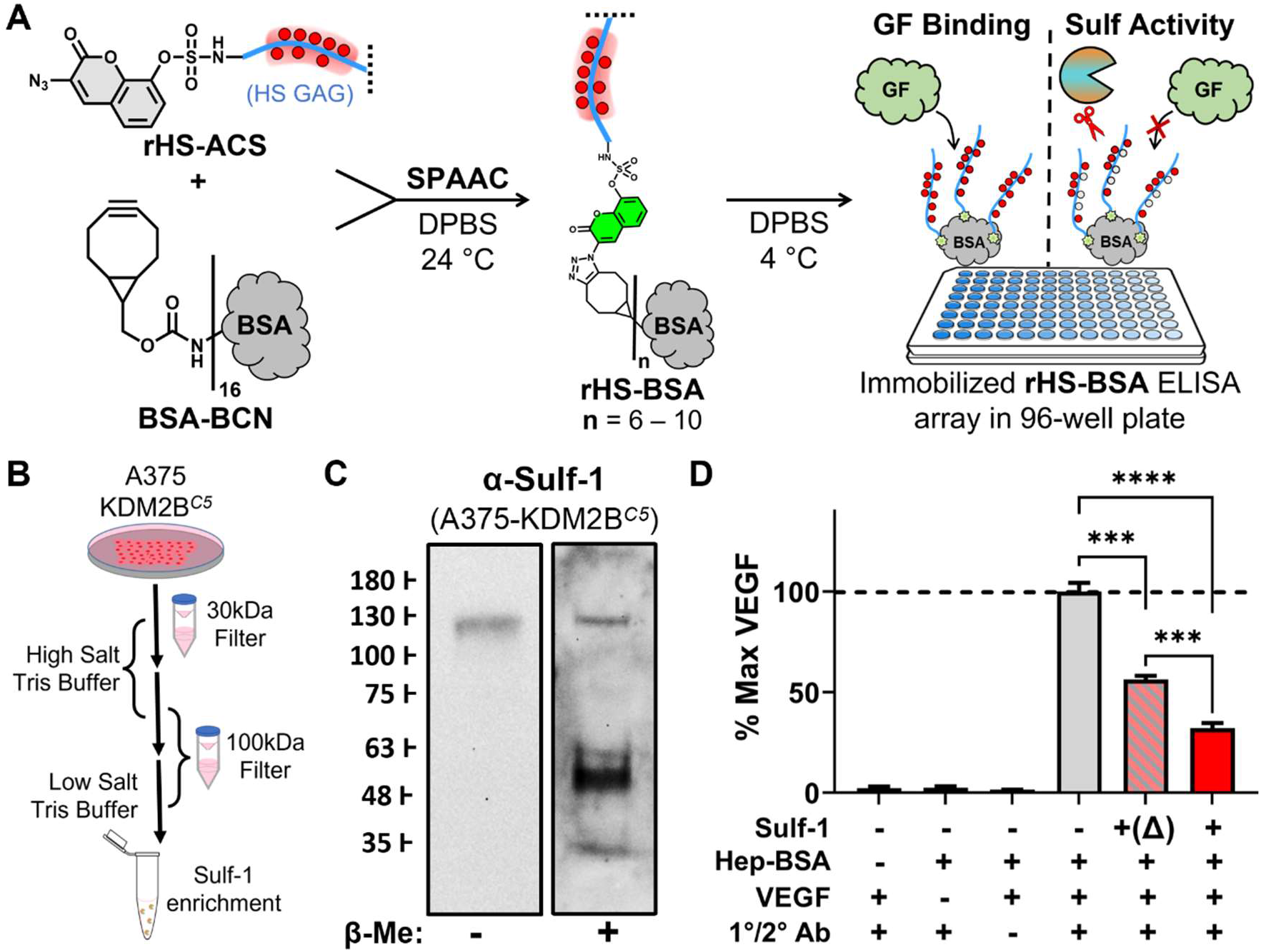
HSPG-mimetic ELISA platform for profiling the effects of Sulf activity on HS-protein interactions. *(A)* A fluorogenic linker strategy for generating HS-bovine serum albumin (BSA) PG-mimetics. The HS-BSA conjugates are immobilized in 96-well plates and used as a substrate to analyze GF binding and Sulf activity via ELISA *(B)* Enrichment protocol for sourcing Sulf-1 from A375 KDM2B^C5^ cell culture. (*C*) Western blot analysis of Sulf-1 enriched samples with a-Sulf-1 Ab (SAB1410410) in non-reducing and reducing conditions. Full protein ∼ 133 kDa. (*D*) Effects of catalytically active Sulf-1 on VEGF binding to **Hep-BSA** via ELISA. Partial rescue of VEGF response through heat inactivation “**+(Δ)**” of Sulf-1 suggested residual binding of catalytically inactive denatured Sulf-1.

The rHS-BSA conjugates were generated by merging rHS chains primed at their reducing end with a reactive fluorogenic linker, 3-azidocoumarin-7-sulfamide (**rHS-ACS**) and BSA functionalized with complementary bicyclo[6.1.0]nonyne groups (**BCN-BSA**) (**Fig. 2*A***). Subsequent strain-promoted azide-alkyne cycloaddition (SPAAC)^39^ converts the azido group in **ACS** into a triazole, linking the rHS chains covalently to the BSA carrier and releasing fluorescence. The resulting light emission (l = 477 nm) is then used to monitor the reaction and establish the composition of the resulting **rHS-BSA** conjugates (***SI Appendix,* Fig. S2**). Conjugates containing heparin (**Hep-BSA**) and murine liver-derived HS (**LivHS-BSA**) were generated as controls to establish continuity with prior studies. The conjugates (1 – 2 ng/μL in DPBS, 100 µL) were arrayed on high-binding 96-well plates for profiling Sulf activity (**Fig. 2*A***).

Due to challenges associated with the purification and stability of recombinant Sulfs,^40^ it is more accessible to source these enzymes for *in vitro* enzymatic assays using cells with naturally elevated levels of enzyme expression (i.e., MCF-7 cells for Sulf-2 production).^9^ We have adapted the A375 *KDM2B^C5^* cell line^30^ as a convenient source of active Sulf-1. To enrich Sulf-1 and remove low-molecular weight protein components, including endogenous GFs and HS fragments, the conditioned media (CM) were subjected to high-salt treatment and size-exclusion filtration (**Fig. 2*B***). Western Blot analysis with a polyclonal antibody confirmed the presence of Sulf-1 (∼130 kDa), which underwent characteristic fragmentation upon treatment with β-mercaptoethanol and reduction of its disulfide bridge within the HD domain (**Fig. 2*C* and *SI Appendix,* Fig. S3*A***). Treatment of soluble heparin by the Sulf-1 containing fraction resulted in reduction of 6-*O*-sulfation based on LC/MS disaccharide analysis, confirming catalytic activity (***SI Appendix,* Fig. S3*C***).

We validated the PG-mimetic platform for analysis of Sulf activity using the well-established binding of the vascular endothelial growth factor (VEGF) to heparin^41^ and its sensitivity to catalytic removal of 6-*O*-sulfates by the Sulfs.^42,34^ Using ELISA, we compared VEGF binding to immobilized **Hep-BSA** with and without treatment with Sulf-1 (**Fig. 2*D***). Accordingly, **Hep-BSA** was adsorbed overnight on 96-well plates and then incubated with the enzyme at 37 °C for four hours. The plates were then treated with VEGF and binding was detected after incubation with a primary anti-VEGF antibody followed by a secondary antibody-HRP conjugate in the presence of the chromogenic substrate, 3,3′,5,5′-tetramethylbenzidine (*TMB*). As anticipated, Sulf-1 treatment reduced VEGF association by ∼68% relative to the control experiment without Sulf-1 addition. Treatment of **Hep-BSA** with heat-denatured enzyme also resulted in ∼ 44% reduction of VEGF binding (**Fig. 2*D***). The high content of positively charged side-chain residues in the HD domain may have resulted in residual background binding to the **Hep-BSA** in the absence of catalytic activity. We suspected that competitive binding of Sulfs to HS substrates may not be limited to denatured enzymes and set out to evaluate whether it can contribute to the regulation of GF interactions.

### Sulf-1 mediates GF interactions via catalytic remodeling and competitive blocking of HS binding sites

There is growing evidence that catalytically inactive splice variants of Sulfs can regulate GF activity in cells.^43, 44^ Presumably, such effects would be brought about through competition for shared binding motifs within the HS chains. Milz *et al.* and related work by the Dierks group^45^ previously explored the interactions of Sulf-1 with its HS substrate and proposed a link between passive contributions of Sulf binding to HS and impairment of growth factor binding.^46^

To test this possibility, we decoupled the binding and catalytic effects of Sulf-1 on GF interactions by treating the immobilized **Hep-BSA** with Sulf-1 at either 4 °C (blue bars) or 37 °C (red bars), respectively (**Fig. 3*A***). As before, we observed ∼ 67% reduction of VEGF binding at 37 °C; however, treatment at 4 °C resulted in only ∼ 33% reduction compared to untreated **Hep-BSA** control. We investigated whether this dual mechanism for regulation of binding activity extends to other classes of GFs. We tested two members of the fibroblast growth factor (FGF) family of HS-dependent GFs (FGF1 and FGF2) as well as their receptor (FGFR2). While FGF1 has been established to be more sensitive than FGF2 to the presence of 6-*O*-sulfates in HS,^47,48^ this modification is required for signal transduction by both proteins.^49^ Accordingly, the binding of FGF1 and FGFR2 to **Hep-BSA** treated with catalytically active Sulf-1 at 37 °C was reduced to a much larger degree than for FGF2 (**Fig. 3*B***). Interestingly, the catalytically attenuated Sulf-1 was an effective competitor for FGF1 binding sites on heparin compared to FGFR2 and was unable to outcompete FGF2 binding (**Fig. 3*B* and *SI Appendix,* Fig. S4*A***). Thus, FGF2 binding is relatively agnostic to the effects of Sulf-1 and consistent with prior observations for Sulf-2, where there was minimal reduction in FGF2 binding even after 24 hours of digestion with Sulf-2.^21^ FGF2-mediated signaling however, can still be impacted by the Sulfs, ostensibly due to the impact on HS contacts with FGFR.^50^ The differential effect on FGF1 or FGF2 binding caused by the catalytically attenuated Sulf-1 may stem from respective sulfation domain size requirements for FGF1/FGF2 recognition,^48^ a lack of overlap between FGF2 and Sulf-1 binding sites on HS, or comparatively higher FGF2 affinity.

**Figure 3.**
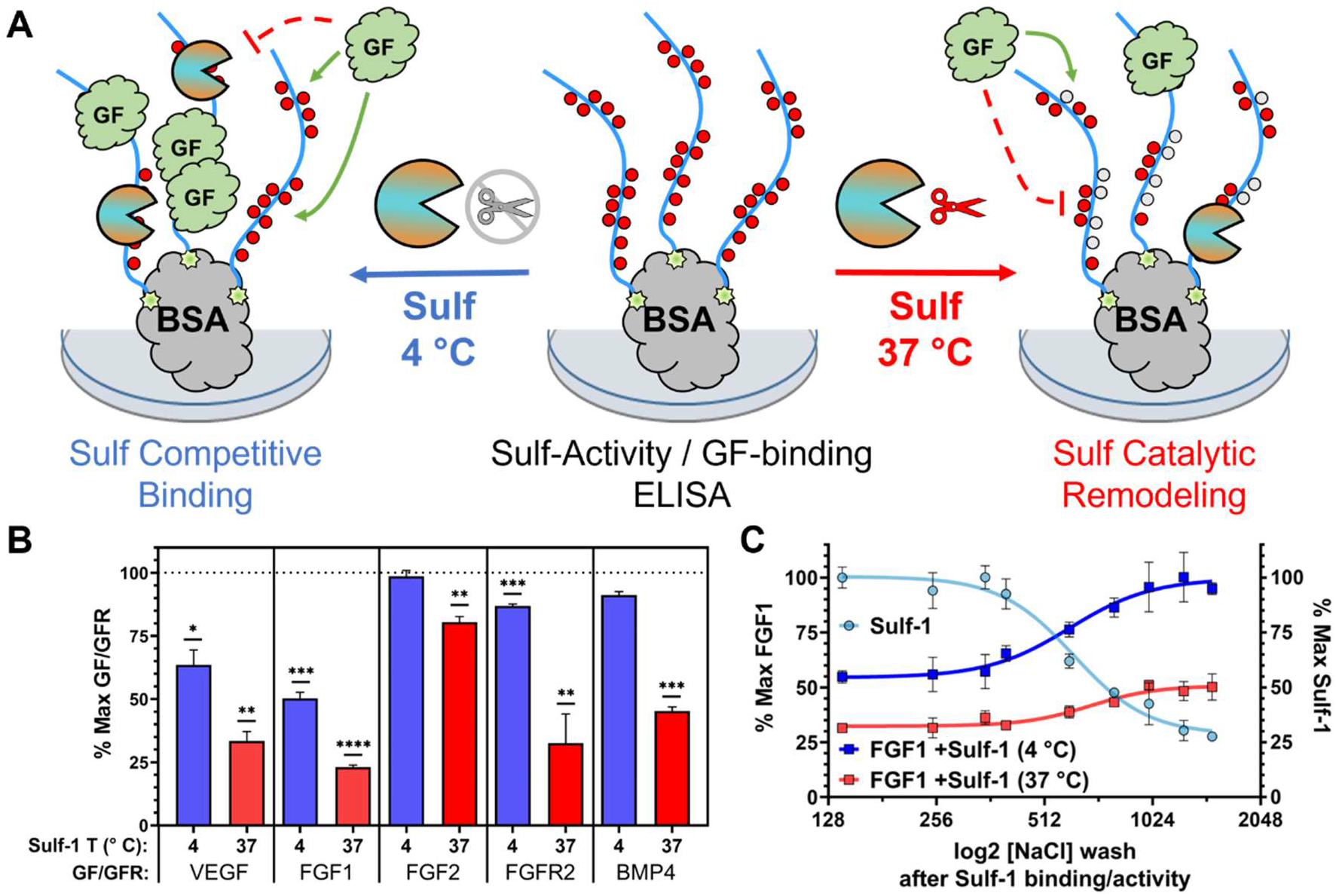
Dual mechanism for regulation of GF-HS interactions by Sulfs. *(A)* A parallel ELISA for analyzing inhibition of GF binding to HS by Sulfs through catalytic processing (37 °C, red) or competitive blocking (4 °C, blue) of substrate binding sites. *(B)* The binding of GFs to **Hep-BSA** after Sulf-1 treatment at 4 °C or 37 °C. Responses are normalized to maximal GF-binding without Sulf-1 treatment at the respective temperature. (*C*) Normalized FGF1 binding to **Hep-BSA** treated with Sulf-1 at 4 °C (blue) or 37 °C (red) and washed with NaCl at increasing concentration. Residual Sulf-1 binding at 4 °C (teal) after salt wash. (Bar graphs and error bars represent mean with SD, n = 3 independent experiments, *p*-values were determined using an unpaired Welch’s t-test, \*\**p* < 0.01, \*\*\**p*< 0.001, \*\*\*\**p*< 0.0001).

In some cases, the effects of Sulf-1 activity on downstream signaling appear context dependent, producing conflicting reports. For instance, prior studies evaluating Sulf regulation of bone morphogenic protein 4 (BMP4) signaling in cells over-expressing the quail ortholog of Sulf-1 (QSulf-1)^51^ or with BMP7 in Sulf-1 mutant mice^50^ showed correlation between signaling and the Sulf expression level, presumably due to the release of Noggin, a BMP antagonist, from HS. The opposite was observed during morphogenesis in zebrafish models.^52^ However, neither study evaluated the effect of Sulf-1 activity on HS-BMP interactions. We tested BMP4 in our assay and observed a ∼ 55% reduction in binding to **Hep-BSA** after treatment with catalytically active Sulf-1 but there was no significant change due to competition from catalytically attenuated Sulf-1, suggesting a possible direct influence of Sulf-1 catalytic activity and 6-*O*-sulfate removal on BMP4 signaling (**Fig. 3*B* and *SI Appendix,* Fig. S4*E***).^41^

To ensure that the observed effects of catalytically attenuated Sulf-1 on GF binding did not stem from residual catalytic activity at 4 °C, we tested whether FGF1 binding could be completely rescued through disruption of Sulf-1 interactions with the Hep-BSA substrate at high salt concentrations (**Fig. 3*C* and *SI Appendix,* Fig. S5*A-B***). Accordingly, we treated **Hep-BSA** with Sulf-1 at either temperature for four hours and the enzymatically treated substrates were then washed with progressively more concentrated buffered saline and probed for GF binding. The binding of FGF1 and an anti-Sulf-1 antibody was then assessed via ELISA. With increasing salt levels, we observed gradual recovery of FGF1 signal accompanied with reciprocal loss of Sulf-1 binding (**Fig. 3*C***). For **Hep-BSA** treated with Sulf-1 at 4 °C, we observed complete rescue of FGF1 binding with high salt washes (>1 M NaCl) compared to an untreated control, in agreement with competitive binding of catalytically inactive enzyme. Similar saline washes of **Hep-BSA** treated with Sulf-1 at 37 °C returned only partial rescue of FGF1 signal, indicating that the overall loss of binding activity stems primarily from catalytic 6-*O*-desulfation by the enzyme, with a minor component resulting from competitive binding to the processed HS (**Fig. 3*C***). FGF1 binding was not affected by exposure of the immobilized **Hep-BSA** to salt alone, confirming that loss of binding was Sulf-1-dependent and not the result of desorption of **Hep-BSA** from the surface (***SI Appendix,* Fig. S5*E***).

### Sulf-1 acts preferentially on highly sulfated HS substrates and the presence of 3-O-sulfate enhances activity

The efficiency of our new conjugation strategy for generating **HS-BSA** conjugates allowed us to expand the scope HS structures beyond heparin to evaluate structure-activity relationships in Sulf-1 regulation of GF interactions. We prepared a set of conjugates comprising rHS chains with varying levels and types of sulfation as well as included HS from mouse liver tissue (**LivHS**) (**Fig. 4*A***). The rHS structures were selected to mirror the levels of sulfation commonly found in native HS structures (e.g., **rHS01**, **rHS02** or **rHS08**) but either lacked or presented a specific sulfation motif (e.g., **rHS02** had minimal 2-*O*-sulfation while **rHS08** presented 3-*O*-sulfates). A substrate, **rHS09**, displaying high levels of sulfation similar to heparin but lacking 3-O-sulfation was also included.

**Figure 4.**
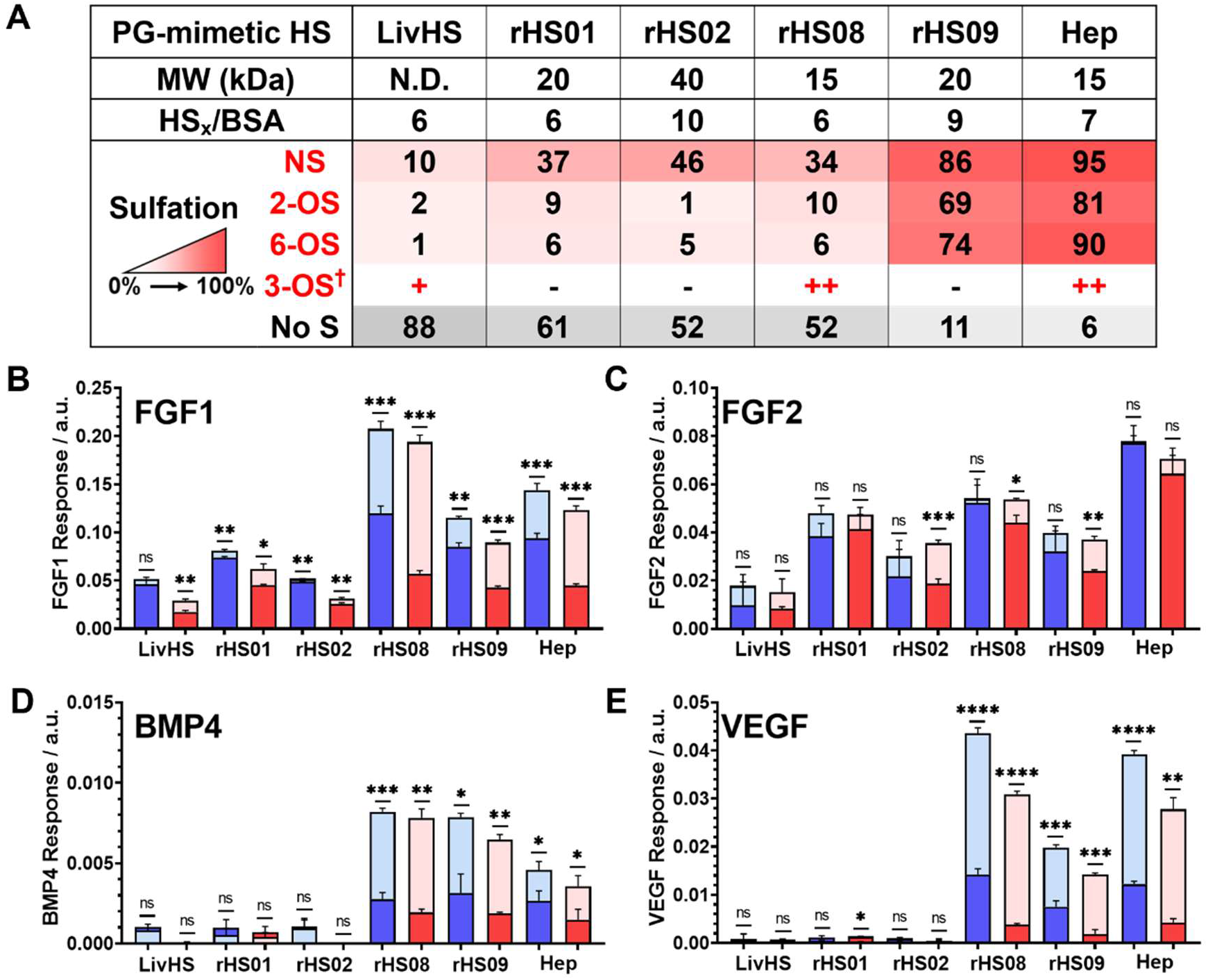
Effects of substrate composition on Sulf-1 regulation of GF binding to HS. *(A)*. Structural characteristics of a panel of PG-mimetics comprising recombinant HS (rHS) structures with defined sulfate compositions for profiling substrate-specificity of Sulfs. Sulfation value and red color intensity indicate % of disaccharides bearing the indicated sulfation type. ^†^3-OS is present based on 3-*O*-sulfotransferases expression, but value could not be determined based on disaccharide analysis. Binding of FGF1 *(B)*, FGF2 *(C),* BMP4 *(D),* and VEGF *(E)* to PG mimetics treated with Sulf-1 at 4 °C (blue) and 37 °C (red). Binding is referenced to maximal responses without Sulf-1 treatment at the respective temperatures of 4 °C (light blue) and 37 °C (light red). (Bar graphs and error bars represent mean with SD, n = 3 independent experiments, *p*-values were determined using an unpaired Welch’s t-test, \*\**p* < 0.01, \*\*\**p*< 0.001, \*\*\*\**p*< 0.0001).

We tested a panel of four GFs (FGF1, FGF2, VEGF and BMP4) against the array of PG-mimetic substrates (**Fig. 4*B-D***) in the presence of Sulf-1 with full (37 °C, red bars) or attenuated (4 °C, blue bars) activity. The binding of GFs to the individual HS-BSA substrates without Sulf1 treatment was used as a reference for maximal binding (**Fig. 4*B-D***, semitransparent bars). To account for any desorption of rHS-BSA structures from the ELISA plates during enzyme incubation, we assessed GF binding at both temperatures.

For FGF1 and FGF2 we observed binding to glycan chains exhibiting both low (**LivHS**, **rHS01**, **rHS02**, **rHS08**), and high (**rHS09** and **Hep**) sulfation, with FGF1 showing increased preference for the latter (**Fig. 4*B* and 4*C***). By contrast, VEGF and BMP4 strongly favored binding to the highly sulfated **Hep** and **rHS09** chains as well as the 3-O-sulfated **rHS08** substrate (**Fig. 4*C* and 4*D***).

Consistent with prior *in vitro* enzymatic assays establishing that Sulf-1 acts most efficiently on *N*-, 2-*O*- and 6-*O*-trisulfated disaccharides in HS oligosaccharide sequences,^27,28^ we observed higher impact of Sulf-1 on the binding of FGF1, VEGF and BMP4 to conjugates with higher proportion of this structural motif (i.e., **rHS09** and **Hep**). Interestingly, in the case of **rHS08**, we observed an increase in the observed effect of active Sulf-1, presumably from the presence of 3-*O*-sulfates, when compared to the analogous **rHS01** structure that bears similar overall sulfation levels but specifically lacks 3-*O*-sulfates. Sulf-1 was able to exert its effects on the interactions of these three proteins with HS when catalytic activity was diminished at 4 °C, suggesting a blocking mechanism of Sulf-1 competitive binding. However, the impact of Sulf-1 on FGF2 binding to the highly sulfated HS substrates, **rHS09** and **Hep**, was limited (**Fig. 4*B***), showing a surprising stronger impact to the low-sulfation **rHS02**. With all four growth factors, a significant response to Sulf-1 activity on **rHS08** was observed, suggesting a correlation between the 3-*O*-sulfate motif and Sulf-1 activity, which may contribute to Sulf-1 regulation of GF binding to forms of HS that may have otherwise low levels of sulfation. Furthermore, experiments that only consider the catalytic effect of the Sulfs or that use only highly-sulfated forms of HS, such as heparin, may leave biologically relevant molecular interactions obscured.

### Sulf-1 and Sulf-2 isoforms show distinct substrate preferences

One of the main ambiguities surrounding the Sulfs is how they exert their isoform-specific activities and, thus, uniquely influence development and disease. Using our rHS PG-mimetic array, we cross-examined the substrate preferences of Sulf-1 against Sulf-2 enriched from a mastocytoma (MST17B10) cell line with Sulf-2 production driven by the EF1a promoter (**Fig. 5**).^37^

**Figure 5.**
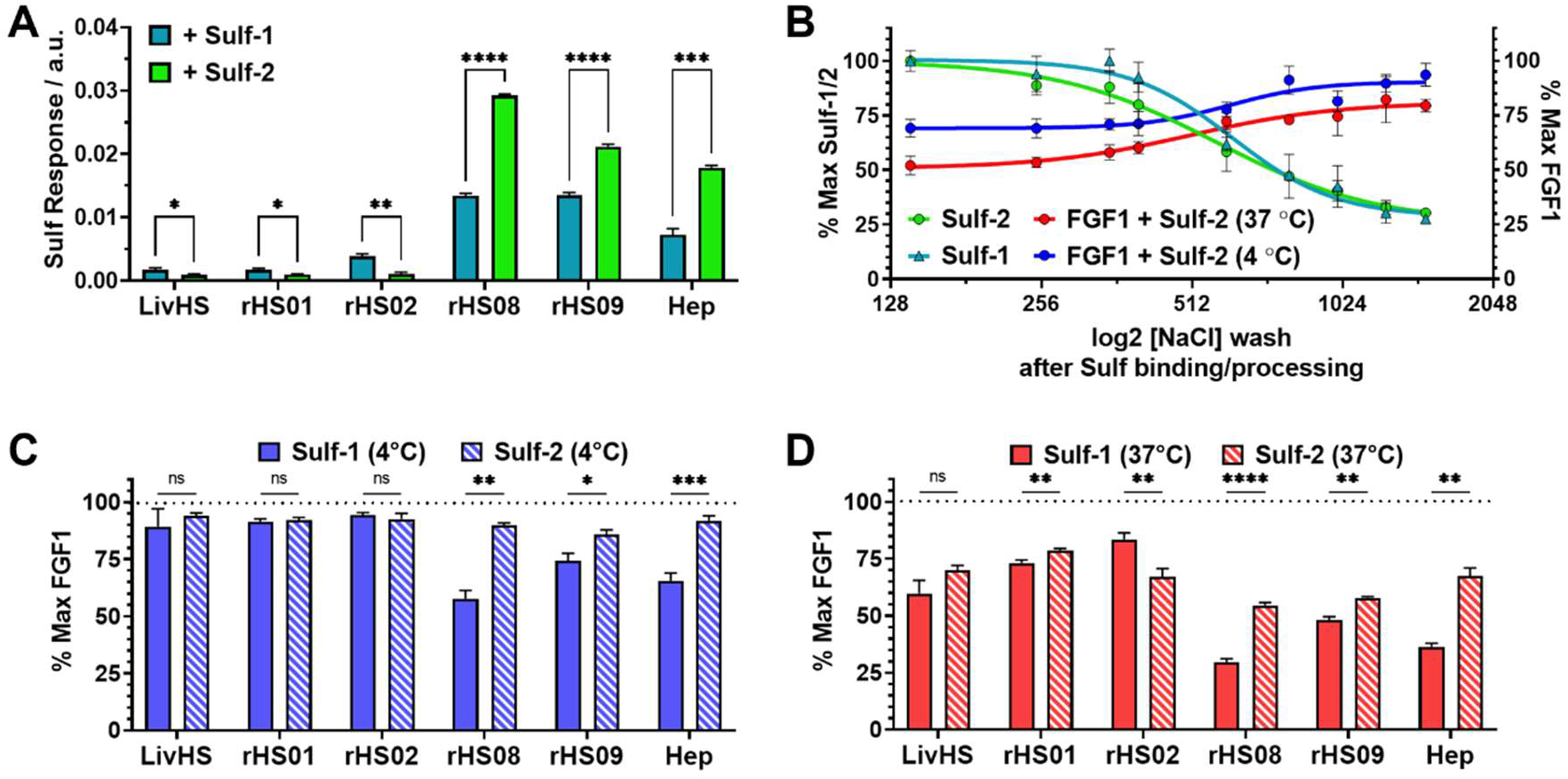
Exploring ELISA-based Sulf-1/2 substrate specificity and activity in the HS-Sulf-FGF1 interactome. *(A)* Sulf-1 and Sulf-2 binding to immobilized rHS-BSA conjugates. *(B)* Residual Sulf-2 binding to Hep-BSA after incubation at 4 °C followed by an NaCl gradient wash (green line). Sulf-1 binding data after similar treatment is included for comparison (teal line). Normalized FGF1 binding to Hep-BSA after reaction with Sulf-2 at 4 °C (blue) and 37 °C (red) and NaCl gradient treatment. *(C)* FGF1 binding to immobilized rHS-BSA conjugates after Sulf-1 and Sulf-2 treatment at 4 °C. FGF1 binding is normalized to maximal binding response in the absence of Sulf (dotted line). (*D*) FGF1 binding to rHS-BSA treated with Sulf-1 and Sulf-2 at 37 °C and washed with NaCl gradient. FGF1 binding is normalized to maximal signal in conditions without Sulf (dotted line). (Bar graphs and error bars represent mean with SD, n = 3 independent experiments, *p*-values were determined using an unpaired Welch’s t-test, \*\**p* < 0.01, \*\*\**p*< 0.001, \*\*\*\**p*< 0.0001).

First, we evaluated the binding of both Sulfs across the panel of rHS conjugates via ELISA (**Fig. 5*A***). The Sulfs were incubated with the immobilized conjugates at 4 °C and detected using isoform-specific antibodies. The binding profile for Sulf-1 mirrored our observations of increased impact of Sulf-1 on GFs interactions with more highly sulfated substrates (i.e., **rHS09** and **Hep**) and the 3-*O*-sulfated HS analog (**rHS08**) (**Fig. 4*D***). The binding of Sulf-2 showed a similar trend; however, the change in binding response between the low- and high-sulfation HS variants was greater compared to Sulf-1 (**Fig 5*A***). In the subset with lower overall sulfation, Sulf-1 showed slightly higher response, compared to Sulf-2, with a significant difference for **rHS02**. This trend was reversed for conjugates with higher levels of sulfation, with Sulf-2 exhibiting significantly higher binding to **rHS08**, **rHS09**, and **Hep** conjugates. Interestingly, examination of the release of both enzymes from immobilized **Hep-BSA** under NaCl gradient followed similar profiles, with half-maximal dissociation for Sulf-1 and Sulf-2 observed at 618 ± 72 mM and 572 ± 75 mM, respectively (**Fig. 5*B***). These values were consistent with those previously reported by Milz *et al*. for the release of catalytically inactive Sulf-1 from porcine intestinal mucosa HS (∼870 mM NaCl) and from 6-*O*-desulfated HS that was processed prior to binding experiments with an active form of Sulf-1 (∼710 mM NaCl).^46^

One possible rationale for these observations is that the isoform-specific binding of Sulfs may not be defined by the large differences in the strength of their electrostatic interactions with preferred HS sequences, but rather by the frequency of these preferred sites within the glycan chain. Sulf-2 appears to favor sequences with higher sulfate content and charge density, which are more prevalent in heparin and the heparin-like substrate, **rHS09**, and present in regions of **rHS08** containing 3-*O*-sulfates.

Since the mechanism by which Sulfs exert effects on GF binding is a composite of competitive binding and catalytic remodeling of HS substrates, differential enzymatic processing abilities may also contribute to isoform-specificity. Therefore, we sought to compare the effects of competitive binding (**Fig. 5*C***) and catalytic activity (**Fig. 5*D***) of Sulf-1 and Sulf-2 on FGF1 interactions with rHS substrates. Consistent with their binding profiles (**Fig. 5*A***), neither enzyme exhibited significant ability to compete with FGF1 for the lower-sulfation subset of **LivHS**, **rHS01**, and **rHS02** substrates at 4 °C (**Fig. 5*C***). In the case of the more sulfated **rHS09** and **Hep** structures as well as the 3-*O*-sulfated **rHS08**, Sulf-1 reduced FGF1 binding to a larger extent than Sulf-2, despite the better binding of the latter to these glycans. This suggests a greater overlap between binding sites for FGF1 and Sulf-1.

With normal enzymatic activity at 37 °C, both enzymes reduced FGF1 binding across the entire set of rHS conjugates (**Fig. 5*D***). Both Sulfs exhibited similar reductions in FGF1 binding within the less sulfated set of substrates (**LivHS**, **rHS01** and **rHS02**), although we observed slightly attenuated Sulf-1 activity on **rHS02**, which contains reduced 2-*O*-sulfate levels. Increasing substrate sulfation correspondingly enhanced FGF1 inhibition by Sulf-1 but, interestingly, did not produce the same response for Sulf-2. Within the higher-sulfation conjugate trio, Sulf-1 exerted a stronger effect on **rHS08** and **Hep**, both of which present 3-*O*-sulfates. The apparent enhancement in processing activity of Sulf-1 is also apparent in the recovery of FGF1 binding enzymatically treated Hep-BSA after high salt wash. While Sulf-1 reduced FGF1 binding by ∼50% due to catalytic desulfation of the substrate (**Fig. 3*C***, red line), the Hep-BSA still retained ∼ 75% of its FGF1 binding after treatment with Sulf-2 (**Fig. 5*B***, red line). The observed activity differences for Sulf-1 and Sulf-2 agree with their relative rates for processing of HS substrates reported by Seffouh *et al*.,^21^ and the higher processing power of Sulf-1 in the PG-mimetic panel appears to be correlated with the presence of 2-*O* and 3-*O*-sulfate modifications within their rHS chains.

### Soluble heparin competitor removes cell surface Sulf-1 and enhances FGF1 binding capacity

To confirm that competitive blocking of shared HS binding sites by Sulfs contributes to the regulation of GF interactions at the cell surface, we assessed changes in the FGF1 binding capacity of endogenous HS structures after Sulf-1 removal using a cell-surface ELISA (**Fig 6**). While convenient for disrupting Sulf-HS interactions in *in vitro* assays, high salt treatment also induces the dissociation of cells from culture substrates. As an alternative, we elected to use increasing amounts of soluble heparin as a competitive ligand to sequester Sulf-1 from cultured A375 KDM2B^C5^ cells.^53^ The treatment resulted in a decrease in cell surface-associated Sulf-1 (**Fig 6A**) and concurrent enhancement of FGF1 binding (**Fig 6B**). The increase in FGF1 binding cannot be attributed solely to sites vacated by Sulf-1, as heparin is likely to release other HS-binding proteins that may share binding motifs with FGF1. The release of cell-surface Sulf-1 is consistent with our findings that there is a significant population of active Sulf-1 bound to HS, using a dual mechanism of catalytic remodeling and competitive binding to regulate HS-GF interactions.

**Figure 6.**
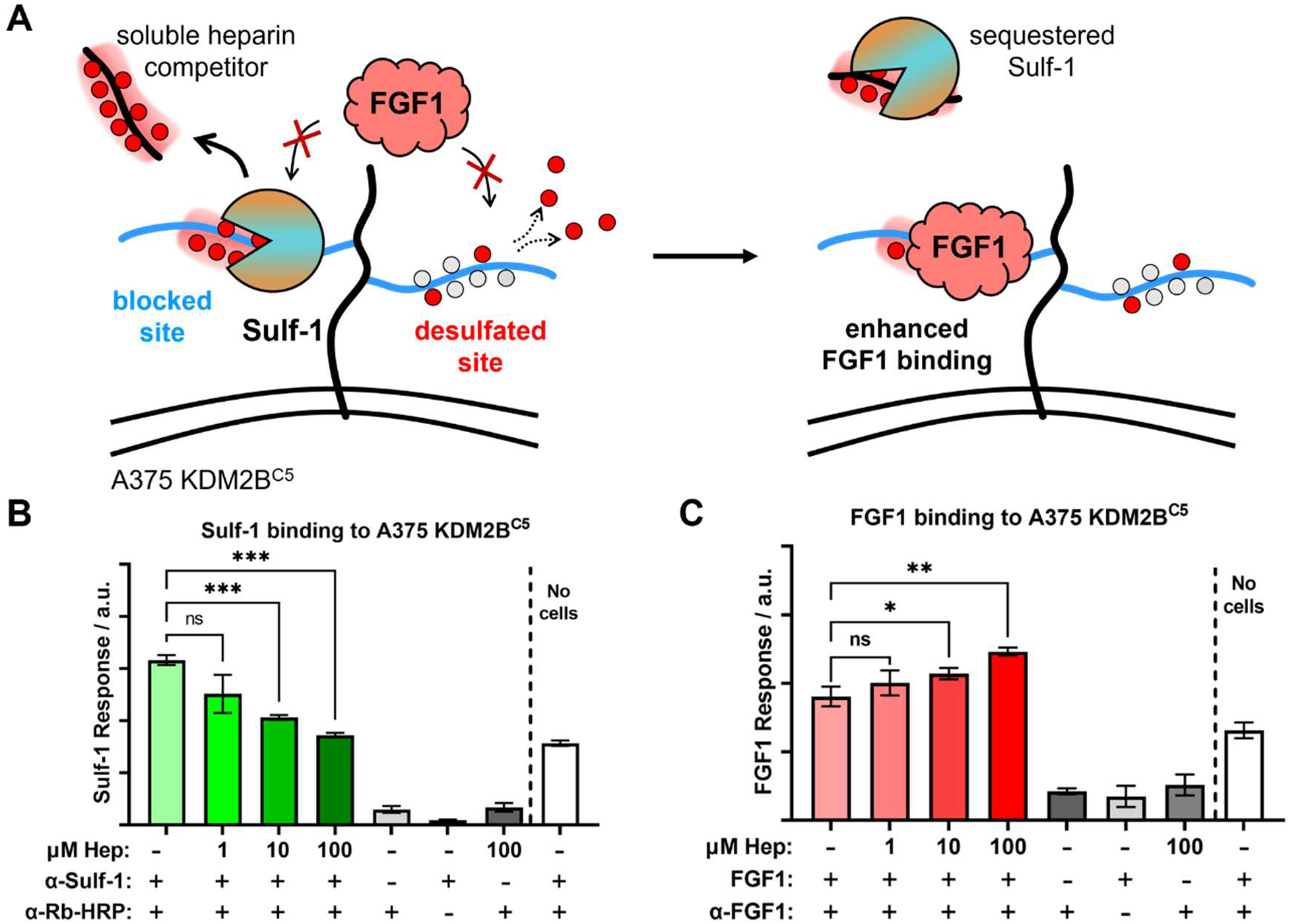
Heparin reduces cell surface-associated Sulf-1 and rescues binding of FGF1 by A375 KDM2B^C5^ cells. (*A*) Soluble heparin acts as a competitive decoy substrate for Sulf-1. Sites in HS vacated after Sulf-1 sequestration become available for FGF1 binding. (*B*) Sulf-1 binding to A375 KDM2B^C5^ cells after washing with increasing amounts of heparin via ELISA. (*C*) FGF1 binding to A375 KDM2B^C5^ cells after washing with increasing amounts of heparin via ELISA. (Bar graphs/points and error bars represent mean with SD, n = 3 independent experiments, *p*-values were determined using a one-way ANOVA (Brown-Forsythe/Welch), \*\**p* < 0.01, \*\*\**p*< 0.001, \*\*\*\**p*< 0.0001).

## Conclusion

In conclusion, we explored the mechanisms by which Sulf-1 and Sulf-2 regulate interactions between cell-surface heparan sulfate and GFs, confirming the known ability of Sulfs to alter the binding specificity of HS by catalytically remodeling its sulfated regions. Notably, we profiled a second mechanistic component, whereby Sulfs act as competitive binders that occupy and block growth factor binding sites. The relative contribution of each mechanism will depend on the HS processing ability of each respective Sulf as well as the overlap between the Sulf and GF binding regions within HS chains and will thus be influenced by the sulfation characteristics of HS. Analysis of these parameters using a panel of engineered rHS conjugates with defined sulfation compositions revealed preferences of Sulf-1 and Sulf-2 for the binding to subsets of HS with low and high levels of sulfation, respectively. Sulf-1 also exhibited increased capacity to catalytically remodel the more highly sulfated HS chains, with 2-O- and 3-O-sulfation motifs possibly driving in the enhanced activity. Sulfs regulation of binding is also GF-specific, with Sulf-1 inhibiting FGF1, VEGF, and BMP4 binding through both competition and catalytic processing and to a greater extent than for FGF2. We anticipate that a deeper examination of these unique relationships between HS structure, Sulf activity and GF binding may help explain the functional differences of the Sulfs observed in biological systems.

## Materials and Methods

### Generation of HS-BSA Conjugates

Additional data for rHS-BSA conjugation, ELISAs, and gels/blots can be found in the SI Appendix, Supplementary Information Text. All chemical and biological reagents and solvents were sourced as indicated in the methods sections and used as received according to manufacturer’s recommendation. rHS glycans and conditioned media from Sulf-2-expressing MST17B10 cells were provided by TEGA therapeutics. Heparin (Iduron), rHS (TEGA), or other GAG disaccharide compositions were either provided in data sheets by the manufacturer or characterized by LC/MS comparison to heparin dp2 standards at the Glycoscience Research and Training Center (UCSD, San Diego).

### Generation of HS-BSA PG mimetic substrates

HS-ACS and BCN-BSA reagents for generating HS-BSA conjugates were synthesized as previously reported.^36^ PBS solutions of the coupling components (50 µL, 1 mM of HS-ACS or ACS-F and 50 µL, 0.5 mg/mL in water of BCN-BSA) were combined in respective wells of a black, clear-bottom 96-well plate and diluted with PBS to a final volume of 200 µL and 1: 6 ACS/BCN ratio. Using a microplate spectrophotometer (Molecular Devices), kinetic fluorescence readings were collected with Ex. 393 nm/ Em. 477 nm for 26 hrs to assess reaction progress. The conjugation of ACS-F to BCN-BSA was determined to occupy all 16 available BCN sites and the fluorescence output was used as a benchmark to determine the conjugation of HS-ACS to BCN-BSA. After this time, the wells were collected and unreacted HS-ACS or ACS-F were removed by spin-filtration (50 kDa MWCO, 5 x 500 µL). The retentates were collected (40 µL), transferred into separate Eppendorf tubes, and diluted with water to a final concentration of 200 ug/mL BSA as determined by BCA assay. The HS-BSA conjugates were further diluted (1 – 2 ng/µL) for adsorption onto 96-well plates for binding assays.

### Cell Culture and Sulf-1 Enrichment

Cells were cultured in monolayer in a T-175 tissue culture flask with DMEM + 10% FBS to a confluency of 80%. The media was replaced with 16 mL fresh Opti-MEM and the cells were incubated for 48 hours, after which 4 mL 5 M NaCl, 50 mM Tris++ buffer (50 mM Tris, 5 mM MgCl_2_, 5 mM CaCl_2_, 5 M NaCl, 1x PI, pH 7.5) was added to the media. The conditioned media was then collected and centrifuged (2000xg, 5 min), and the supernatant was passed through a 0.2 μM nylon filter. The filtered media was transferred to a 30 kD MWCO polyethersulfone spin filter (Thermo Scientific-Pierce). The sample was concentrated by centrifugation (4200xg, 30 min), and the media was exchanged three times by the addition of 50 mM Tris++ (high salt) buffer (50 mM Tris, 5 mM MgCl_2_, 5 mM CaCl_2_, 1 M NaCl, 1x PI, pH 7.5) to the concentrated sample, centrifuging to concentrate between washes. The 30 kDa filtered supernatant was transferred to a 100 kDa MWCO polyethersulfone spin filter (Thermo Scientific-Pierce). The sample was concentrated by centrifugation (4200xg, 30 min), and the media was exchanged three times by the addition of 50 mM Tris++ (low salt) buffer (50 mM Tris, 5 mM MgCl_2_, 5 mM CaCl_2_, 100 mM NaCl, 1x PI, pH 7.5) to the concentrated sample, centrifuging to concentrate between washes. The resulting solution was quantified for total protein content using the BCA Assay (Thermo Scientific-Pierce) and then transferred to an Eppendorf tube charged with equal volume 66% (v/v) high purity glycerol / Tris++ buffer, supplemented with PI (1X), aliquoted, and stored at -20 °C.

### ELISA Protocols

All ELISA assays were conducted in high-binding 96-well plates (Greiner 655097), following a general set-up protocol. In triplicate, sample wells were charged with a 100 µL of purified Heparin-BSA or rHS-BSA solution (1 – 2 ng/µL) in DPBS++ (Gibco), and the plates were rocked overnight (16 – 24 hours) at 4 °C. The wells were then thrice-washed with DPBS++, and blocked with 2% BSA:DPBS++ for 1 hour at rt. The plates were then processed according to the intended assay. (See below). After all enzymatic reactions and binding events, the wells were twice-washed with 0.1% Tween-20 (v/v) DPBS++, once with DPBS++, and then charged with TMB substrate solution (100 µL per well, VWR). (For experiments involving NaCl gradients, wells received an additional 10 min incubation with NaCl-supplemented DPBS++ after the first wash.) Immediately after, the kinetic absorbance at 370nm (10 min, 30 – 60s interval) was measured in a SpectraMax i3x plate reader (Molecular Devices), and the results plotted in Prism 9.0 (Graphpad). Results are representative of three or more separate experiments, evaluating the linear range of signal response in each sample.

### Sulf Activity / Growth Factor ELISA

After HS-BSA binding and blocking steps, wells were washed with DPBS++, Sulf samples in Tris++ buffer (50 mM Tris, 5 mM MgCl_2_, 5 mM CaCl_2_, pH 7.5 – 0.0.1 µg/µL total protein) were added to non-control wells, while non-treated wells received 0.2% BSA:Tris++. For tandem temperature ELISAs, one of two duplicate plates was incubated at 37 °C for 4 hours, and one plate at 4 °C. All wells were then twice-washed with 0.1% Tween-20 (v/v) DPBS++, once with DPBS++, and then blocked with 2% BSA:DPBS++ for 1 hour at rt. All wells were then twice-washed with 0.1% Tween-20 (v/v) DPBS++, then once with DPBS++, and recombinant growth factor was added (100 µL, 10 - 50 nM in 0.2% BSA:DPBS++). The plates were then incubated overnight (12 – 16 hours) at 4 °C. The next day, the wells were twice-washed with 0.1% Tween-20 (v/v) DPBS++, once with DPBS++, and then supplemented with the corresponding pre-complexed primary and secondary-HRP antibodies (100 µL in 0.2% BSA:DPBS++) and rocked for 1.5 hours at room temperature. Wells were twice-washed with 0.1% Tween-20 (v/v) DPBS++, once with DPBS++, and then analyzed with addition of TMB substrate according to the general protocol above.

### Sulf Binding ELISA

Sulf samples in Tris++ buffer (50 mM Tris, 5 mM MgCl_2_, 5 mM CaCl_2_, pH 7.5 – 0.0.1 µg/µL total protein) were pre-complexed with corresponding primary antibodies for 6 hours at 4 °C. The samples were then added to 96-well plates after HS-BSA binding and blocking steps as indicated in the general ELISA protocol, and incubated for 12 hours at 4 °C. The next day, the wells were twice-washed with 0.1% Tween-20 (v/v) DPBS++, once with DPBS++, and then supplemented with corresponding secondary-HRP antibodies (100 µL in 0.2% BSA:DPBS++) and rocked for 1.5 hours at room temperature. Wells were twice-washed with 0.1% Tween-20 (v/v) DPBS++, once with DPBS++, and then analyzed with addition of TMB substrate according to the general protocol above.

### SDS-PAGE and Western Blot Analysis of Sulfs

Aliquots of concentrated C.M. were prepared with matching protein levels, analyzed by SDS-PAGE (4-15% TGX Mini Protean Gel, BioRad), and transferred onto PVDF membranes using standard Western Blotting protocols. Total protein content in the SDS-PAGE gel was visualized by washing the gel in InstantBlue Coomassie Protein Stain (Expedeon/Abcam) overnight at 4 °C. The membranes were processed by blocking with 3% BSA:TBST for one hour at rt, followed by incubation with anti-Sulf-1/2 primary antibody (anti-Sulf-1: rabbit polyclonal, Sigma, SAB1410410, anti-Sulf-2: rabbit monoclonal clone 2B4, Sigma, MABC584) overnight at 4 °C. They were then washed three times with 1X TBST, and incubated with a respective secondary HRP-conjugated antibody (anti-rabbit: Cell Signaling Technologies, #7074S, anti-mouse: Cell Signaling Technologies, #7076) for one hour at room temperature. The membrane was then washed three times with TBST, and visualized by the addition of Immobilon Crescendo Western HRP substrate (Millipore) followed by exposure in a Bio-Rad ChemiDoc XRS imaging system (BioRad).

### Cell-Surface ELISA

(Unless otherwise indicated, all steps used volumes of 200 μL) A375 cells were seeded at a density of 25,000 cells/well in a tissue culture treated 96-well plate two days prior to the ELISA experiment and incubated at 37 °C with 5% CO_2_. Then, the media was carefully aspirated and the cells were gently washed twice with PBS. Then, the cells were respectively incubated with PBS supplemented with heparin at the indicated concentrations for 5 minutes at room temperature. The solution was carefully aspirated and the cells were again gently washed twice with PBS. The cells were then fixed by incubating with 4% paraformaldehyde in PBS for 20 minutes at room temperature, and were then gently washed thrice with PBS. The cells were then incubated with 0.1 M glycine in PBS for 30 minutes at room temperature to neutralize residual fixative, and were again washed thrice with PBS. The wells were then blocked by incubating with 5% BSA (w/v) in PBS for 30 minutes at room temperature. After aspirating the blocking solution, the wells for the FGF1 experiment were incubated with FGF1 (50 nM) in 0.5% (w/v) BSA:PBS for 30 minutes at room temperature. For Sulf-1 analysis, no growth factor was added, and instead proceeded directly to addition of antibodies. The plates were then washed thrice with PBS and incubated with primary antibodies (FGF1: 1:500, Sulf-1: 1:100) in 0.5% (w/v) BSA:PBS for 60 minutes at room temperature. The plates were then washed thrice with PBS and incubated with secondary antibodies (anti-rabbit IgG=HRP, FGF1: 1:1000, Sulf-1: 1:500) in 0.5% (w/v) BSA:PBS for 60 minutes at room temperature. The plates were then washed thrice with PBS, and then 150 uL TMB substrate was added to each well. Immediately after, the kinetic absorbance at 370nm (10 minutes, 30 – 60s interval) was measured in a SpectraMax i3x plate reader (Molecular Devices), and the results plotted in Prism 9.0 (Graphpad). Results are representative of three or more separate experiments, evaluating the linear range of signal response in each sample.

## Acknowledgments

This work was supported by a grant from the National Science Foundation (#2004243). KG was supported by the Alfred P. Sloan Foundation (FG-2017-9094) and the Research Corporation for Science Advancement via the Cottrell Scholar Award (#24119). BMT was supported by the Alfred P. Sloan Foundation Minority Graduate Scholarship Program and the UCSD Molecular Biophysics Training Program (T32, GM008326). The authors wish to thank Professor Mia Huang for assistance with sulfatase assay development, Professor Jeffrey Esko for providing A375 KDM2B^C5^ cells, Professor Ryan Weiss for assistance with the Sulf-1 production, and Professor Philip Gordts for useful discussions and thoughtful insights. The authors also thank the UCSD GlycoAnalytics Core and its director, Dr. Biswa Choudhury, for help with glycan analysis.

## Notes

### Competing Interest Statement

The authors have declared no competing interest.

