## Supporting Information for "Human extracellular sulfatases use a dual mechanism for regulation of growth factor interactions with heparan sulfate proteoglycans"

**Supporting Information Text**

**Abbreviations**

**ACS-F =** 3-azidocoumarin 7-sulfonyl fluoride
**BCA =** bicinchoninic acid
**BCN =** bicyclo[6.1.0]nonyne
**BMP =** bone morphogenic protein
**BSA =** bovine serum albumin
**DCM =** dichloromethane
**DMSO =** dimethyl sulfoxide
**ELISA =** enzyme-linked immunosorbent assay
**ESI-MS =** electrospray ionization mass spectrometry
**EtOAc =** ethyl acetate
**FGF =** fibroblast growth factor
**HCl =** hydrochloric acid
**Hep =** heparin
**Hex =** n-hexanes
**HRP =** horse radish peroxidase
**HS =** heparan sulfate
**IR =** infrared spectroscopy
**MALDI-TOF =** matrix-assisted laser desorption/ionization – time of flight
**MQ =** milli-Q ultrapure water
**MWCO =** molecular weight cut-off
**NHS =** N-hydroxysuccinimide
**NMR =** nuclear magnetic resonance
**PBS =** phosphate buffered saline
**PD-10 =** prepacked sephadex® G-25 medium disposable column
**RBF =** round-bottom flask
**SuFEx =** Sulfur (IV) fluoride exchange
**TCS** = triazole coumarin sulfonyl
**TFA =** trifluoroacetic acid
**VEGF =** vascular endothelial growth factor

**Materials and Methods**

All chemical and biological reagents and solvents were sourced as indicated in the methods sections and used as received according to manufacturer's recommendation. Absorbance and fluorescence values for 96-well plate applications were collected on a SpectraMax i3x plate reader (Molecular Devices) with SoftMax Pro software. The data was exported and analyzed in Microsoft Excel 2016 and GraphPad Prism 9 software. Nuclear magnetic resonance (NMR) spectra were collected on a JOEL ECA 500MHz NMR spectrometer. Spectra are reported in parts per million (ppm) on the δ scale relative to the residual solvent as an internal standard. Data are reported as follows: chemical shift (s = singlet, d = doublet, dd = doublet of doublets, t = triplet, q = quartet, br = broad, m = multiplet), coupling constants (Hz), and integration.

Heparin (Iduron), rHS (TEGA), or other GAG disaccharide compositions were either provided in data sheets by the manufacturer or characterized by LC/MS comparison to heparin dp2 standards at the Glycoscience Research and Training Center (UCSD, San Diego).

**Synthesis of 3-azidocoumarin-7-sulfonyl fluoride**

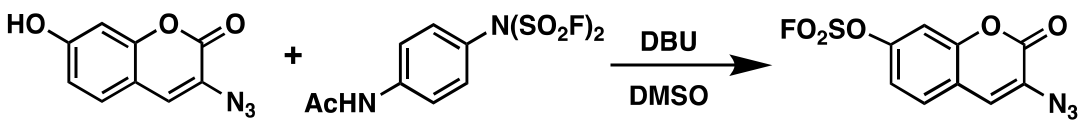

3-azidocoumarin-7-sulfonyl fluoride was prepared as previously described with minor modifications.[i] Briefly, 3-azido-7-hydroxycoumarin (0.1483 g, 0.7299 mmol, 1.0 eq.) and 4-[(Acetylamino)phenyl]imidodisulfuryl difluoride (AISF) (0.2523 g, 0.8028 mmol. 1.2 eq.) were added to a scintillation vial and dissolved in DMSO (4 mL). The vial was flushed with gaseous nitrogen and 1,8-diazabicyclo [5.4.0]undec-7-ene (DBU, 161 µL, 1.08 mmol, 2.2 eq.) was added dropwise. The reaction stirred at room temperature covered in foil for 10 min. The dark brown mixture was diluted in EtOAc and extracted with 0.5 M HCl (2 x 10 mL) then brine (2 x 10 mL). The organic phase was died with anhydrous $MgSO_4$, filtered, then concentrated. The crude brown product was dissolved in EtOAc with trace amount of DCM. Celite was added and the suspension was concentrated and dried under reduced pressure. The crude sample was purified by flash silica chromatography using a linear 10 - 20% EtOAc/Hexane with 0.1% $Et_3N$. Sample fractions were collected, concentrated, transferred to a tared amber vial, and then dried under reduced pressure. A white crystalline solid (0.1605 g, 77%) was obtained. $^1$H NMR (500 MHz, $CDCl_3$) δ 7.54 (d, *J* = 8.6 Hz, 1H), 7.37 (d, *J* = 2.3 Hz, 1H), 7.30 (ddd, *J* = 8.6, 2.4, 0.7 Hz, 1H), 7.21 (s, 1H). $^{13}$C NMR (126 MHz, $CDCl_3$) δ 156.47, 151.52, 149.88, 128.93, 127.91, 124.00, 119.78, 118.10, 110.09 ppm. $^{19}$F NMR: (282 MHz, $CDCl_3$) δ 39.0 (s, 1F).

**GAG conjugation to ACS-F**

Commercial GAGs, including heparin (20 mg, Iduron, Macclesfield SK10 4TG, UK), TEGA recombinant HS (rHS), or Mouse Liver HS, were transferred to a PCR tube and dissolved in 90 µL reaction buffer (1.0 M urea, 1.0 M sodium acetate, pH 4.5). To this solution was added n-methylaminooxy-propylamine linker[ii] (11.5 µmoles). Reducing end conjugation proceeded at 50°C for 24-48 h, after which the reaction was quenched with 200 µL of 2 M Tris-HCl, pH 8.1. The resulting GAG-amine was purified by PD-10 column, followed by concentration and removal of excess linker using 3 kDa molecular weight spin filters (Amicon, Millipore Sigma). To 400 µL of recovered GAG-amine was added 200 µL of 100 mM sodium phosphate, pH 8.0 and 600 µL DMSO. ACSF (38 mg, 133 µmoles) was dissolved in 400 µL DMSO and transferred to the GAG-amine solution. Sulfonyl fluoride exchange (SuFEx) proceeded at ambient temperature for 24 h. The reaction was then diluted with 900 µL water and purified via PD-10 column into a clear 96-well plate. Sample-containing wells were detected by microplate absorbance at 326 nm and were combined and concentrated by 3 kDa spin filtration, followed by lyophilization to afford the desired GAG-ACS.

**BCN-BSA synthesis**

To a 1.5 mL microcentrifuge tube (Fisher Scientific 05408129) was added 1 mL of 100 mM sodium phosphate buffer, pH 8.0, 10 mg BSA (VWR 0332-25G), and 78.1 µL of a 10 mg/mL (1R,8S,9s)-

Bicyclo[6.1.0]non-4-yn-9-ylmethyl N-succinimidyl carbonate (BCN, 17 eq.) (Sigma Aldrich 744867-10MG) and stirred at 4°C for 16 h. The reaction was dialyzed against pure water in 25 kDa molecular weight cut-off dialysis tubing (Spectra 132126), for 48 hours, replacing the water after 24 hours. Lyophilization of the dialyzed product affords 11 mg of the product (quantitative yield). MALDI-TOF MS analysis indicates the modified BSA protein has a molecular weight of about 69,689 daltons compared a starting mass of 66,808 daltons for unmodified BSA.[1] Each additional BCN adds 177.3 daltons, a difference of 3,259 daltons indicates approximately 16 BCN/BSA.

HS-BSA conjugation was then performed as described in the main text.

**Disaccharide Analysis**

For disaccharide analysis, lyophilized HS or Sulf-1 treated Hep was incubated with 2 mU each of heparin lyases I, II, and III for 16 hr at 37 °C in buffer containing 40 mM ammonium acetate and 3.3 mM calcium acetate, pH 7. HS disaccharides were aniline-tagged and analyzed by RP-LC-MS on a LTQ XL Orbitrap mass spectrometer as previously described.[iii] For Sulf-1 activity assays, 5 mg heparin (Iduron) was treated with Sulf-1 (0.1 µg/µL) for four hours at 37 °C in 50 mM Tris, 5 mM MgCl2, 5mM CaCl2, pH 7.5.

**A**

| Sulf 1/2 Physical Characteristics | Catalytic Domain | Hydrophilic Domain | C-Tail Domain | Sulf1/2 Sequence Homology ← |
|---|---|---|---|---|
| | 81% | 45% | 71% | |
| Full-Length MW | N-terminal Subunit | | C-terminal Subunit | PTM |
| 133 kDa | 59.3 kDa | | 38.8 KDa | 34.9 kDa |

**B**

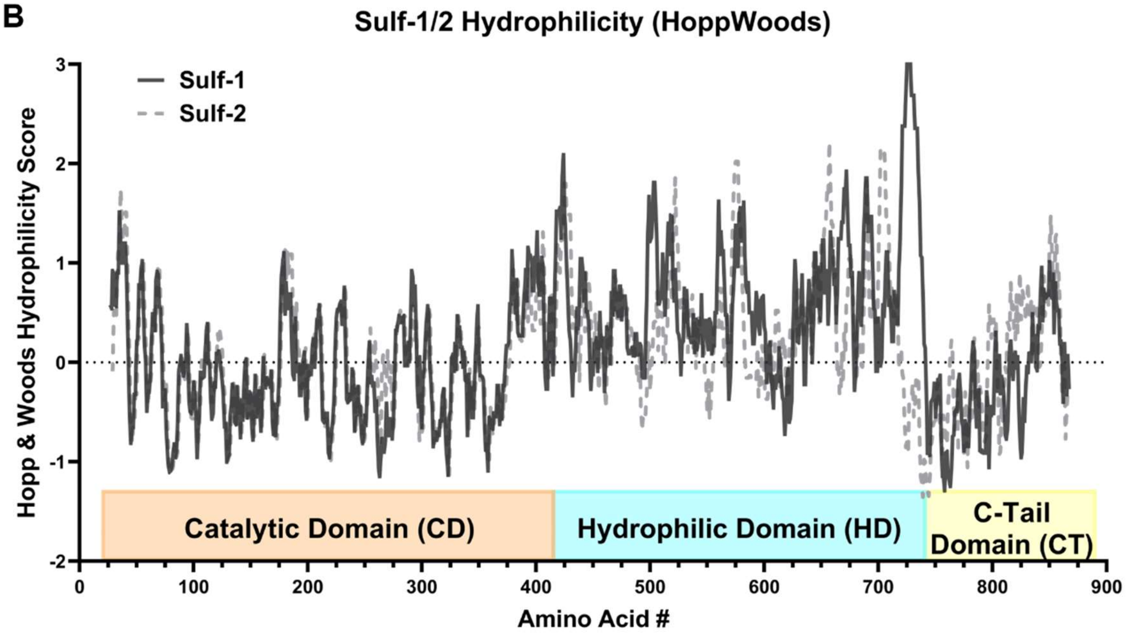

**Fig. S1. Comparison of Sulf-1 and Sulf-2 physical characteristics.** *(A)* Sulf-1 and Sulf-2 sequence homology as determined by BLAST alignment and molecular weight of subunits and posttranslational modifications.[iv] *(B)* Computed hydrophilicity score of Sulf-1 and Sulf-2 (Hopp & Woods)[v] full sequence and *(C)* the hydrophilic-domain.

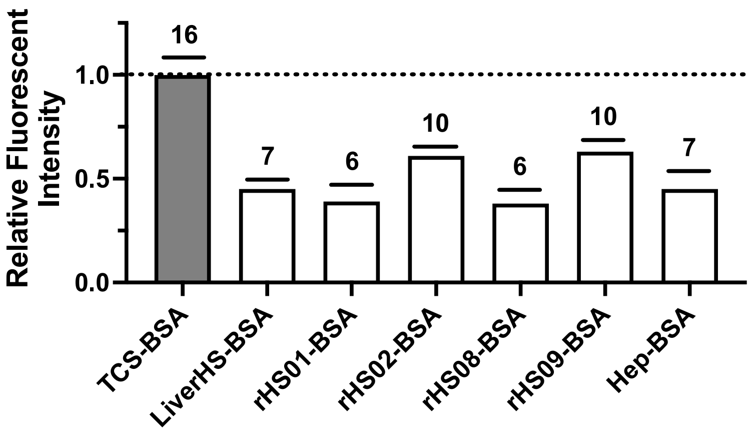

**Fig. S2. Valency of HS-BSA conjugates.** Maximum possible valency of 16 is based on the number of BCN residues per BSA conjugate as determined by MALDI-MS in Porell *et al.*. [1]

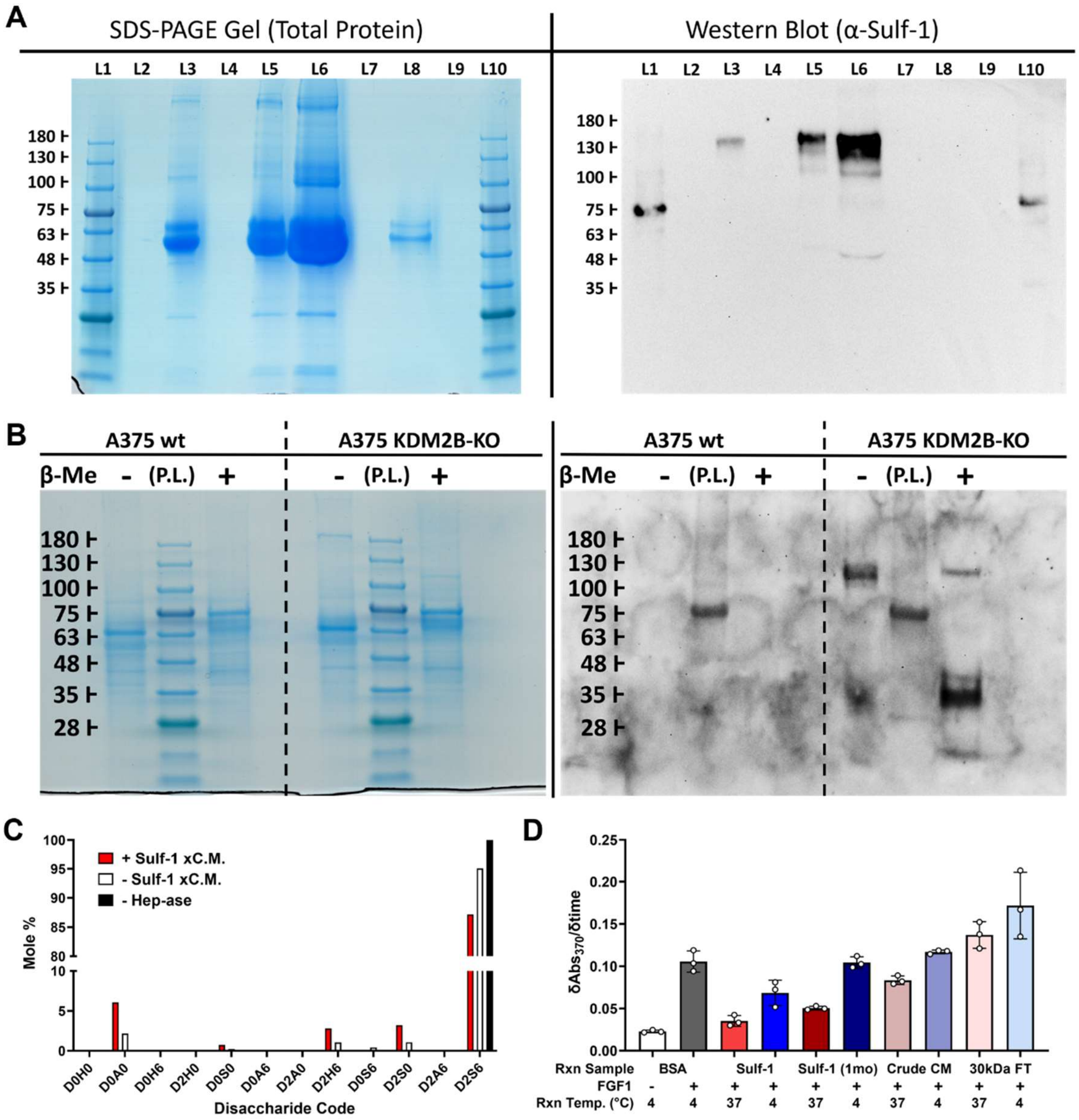

**Fig. S3. Sourcing Sulf-1 from A375 KDM2B$^{C5}$ conditioned media.** *(A)* SDS-PAGE gel (left) of aliquots taken during the Sulf-1 enrichment protocol and corresponding Western Blot (right) stained with a polyclonal Sulf-1 antibody. L1: Protein ladder, L2: Blank, L3: 100kDa-conc. C.M., L4: Blank, L5: Crude C.M., L6: 30kDa-conc C.M., L7: 30kDa filtrate, L8: 100kDa filtrate, L9: Blank, L10: Protein ladder *(B)* SDS-PAGE gel (left) on Sulf-1 enriched media from A375 wt (included for comparison) and A375 KDM2B-KO, loaded with equal protein as normalized by BCA assay, stained with Instant Blue and corresponding Western Blot (right) stained with a polyclonal Sulf-1 antibody. (*C*) LC/MS disaccharide analysis of heparin processed by Sulf-1 enriched conditioned media (red), compared to non Sulf-1 treated heparin (white), and Sulf-1 treated heparin that does not go heparinase digestion (black). (*D*) FGF1 ELISA after reaction with a BSA control, fresh Sulf-1 samples, Sulf-1 stored for 1 month at -20 °C, crude conditioned media, and the resulting flow through after 30 kDa filtration of A375 KDM2B-KO conditioned media.

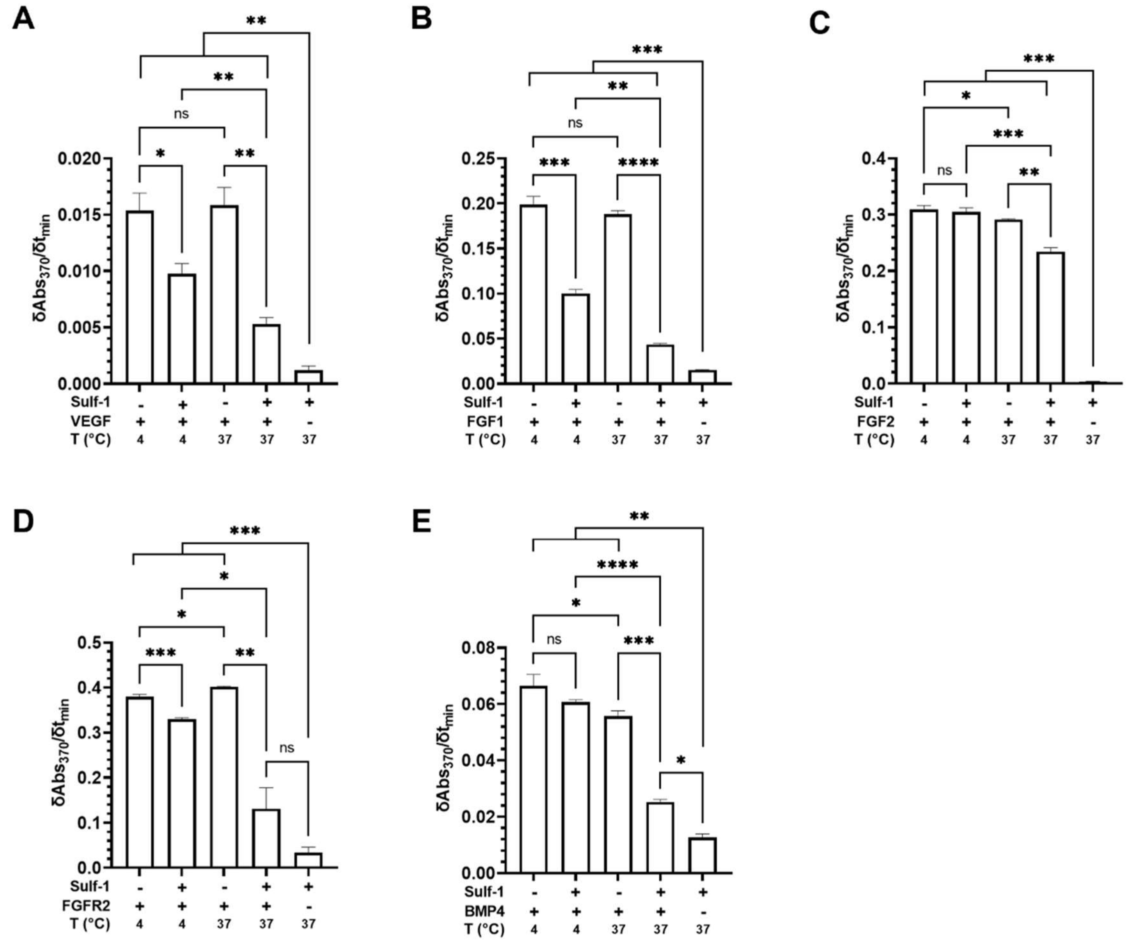

**Fig. S4. Effects of Sulf-1 on GF binding to Hep-BSA.** (Extended data for Figure data for **Figure 3B**). Raw data for the normalized comparison in **Figure 3B**. All samples used 100 uL volume, 200 ng/well **Hep-BSA**, 0.1 ug/uL Sulf-1 xCM. (*A*) 25nM VEGF, 1:500 α-VEGF, 1:1000 α-Rb HRP-IgG. (*B*) 50nM FGF1, 1:500 α-FGF1, 1:1000 α-Rb HRP-IgG. (*C*) 50nM FGF2, 1:500 α-FGF2, 1:1000 α-Ms HRP-IgG. (*D*) 50nM FGFR2, 1:500 α-FGFR2, 1:1000 α-Hu HRP-IgG. (*E*) 50nM BMP4, 1:500 α-BMP4, 1:1000 α-Ms HRP-IgG. (Bar graphs and error bars represent mean with SD, n = 3 independent experiments, *p*-values were determined using an unpaired Welch's t-test, $^{**}p < 0.01$, $^{***}p < 0.001$, $^{****}p < 0.0001$).

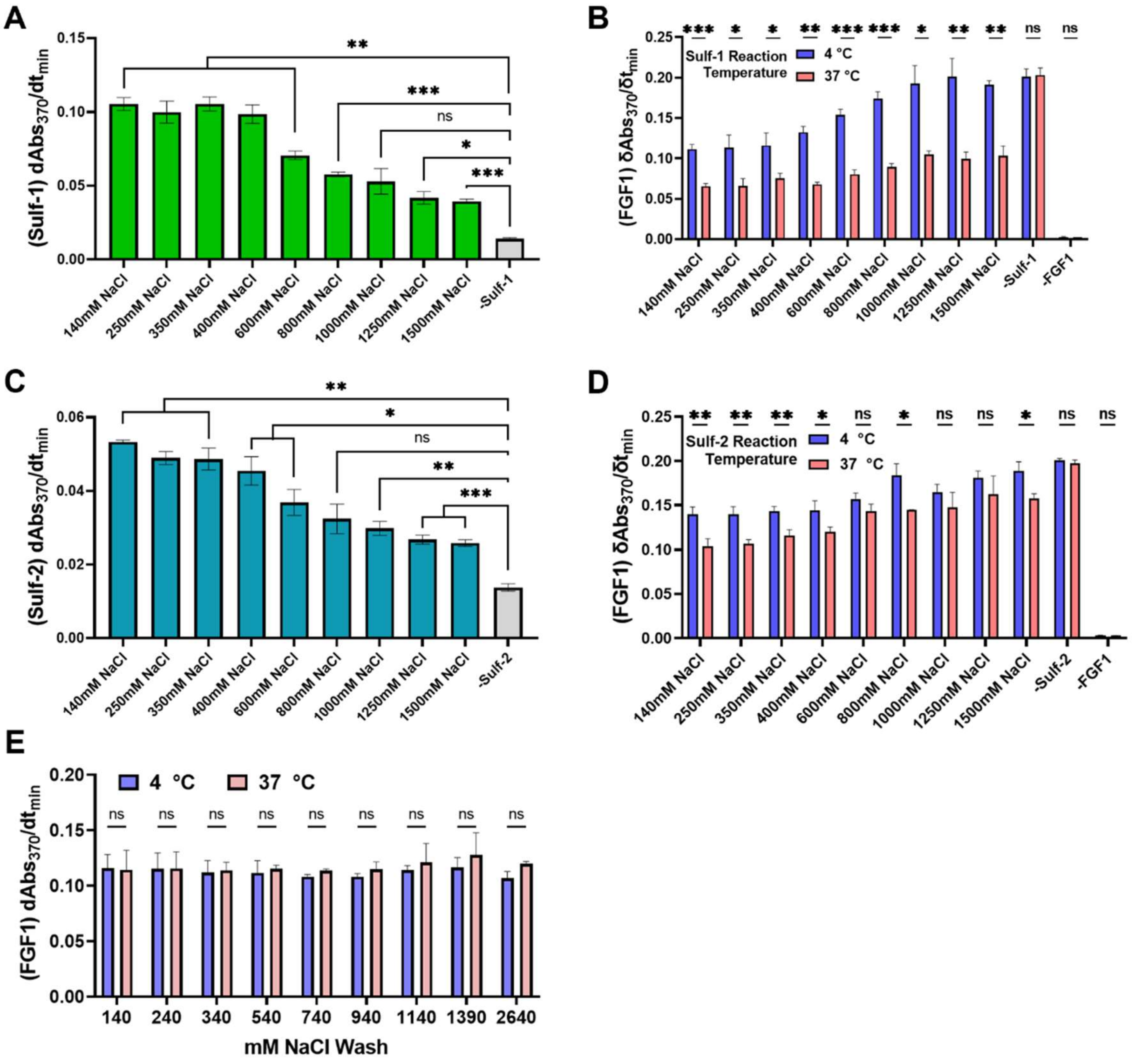

**Fig. S5. Binding of Sulfs and FGF1 to Hep-BSA treated with Sulfs and washed with increasing NaCl concentration via ELISA.** (Extended data for Figures 3C and 5B). Raw data and the stability of immobilized **Hep-BSA** to salt in the absence of Sulf treatment. All samples used 100 uL volume, 200 ng/well rHS$_x$-BSA. (*A*) 0.1 ug/uL Sulf-1 xCM, 1:100 α-Sulf-1, 1:250 α-Rb HRP-IgG. (*B*) 0.1 ug/uL Sulf-1 xCM, 50nM FGF1, 1:500 α-FGF1, 1:1000 α-Rb HRP-IgG. (*C*) 0.1 ug/uL Sulf-2 xCM, 1:100 α-Sulf-2, 1:250 α-Ms HRP-IgG. (*D*) 0.1 ug/uL Sulf-2 xCM, 50nM FGF1, 1:500 α-FGF1, 1:1000 α-Rb HRP-IgG. (*E*) 50nM FGF1, 1:500 α-FGF1, 1:1000 α-Rb HRP-IgG. (Bar graphs and error bars represent mean with SD, n = 3 independent experiments, *p*-values were determined using an unpaired Welch's t-test, ***p* < 0.01, ****p*< 0.001, *****p*< 0.0001).

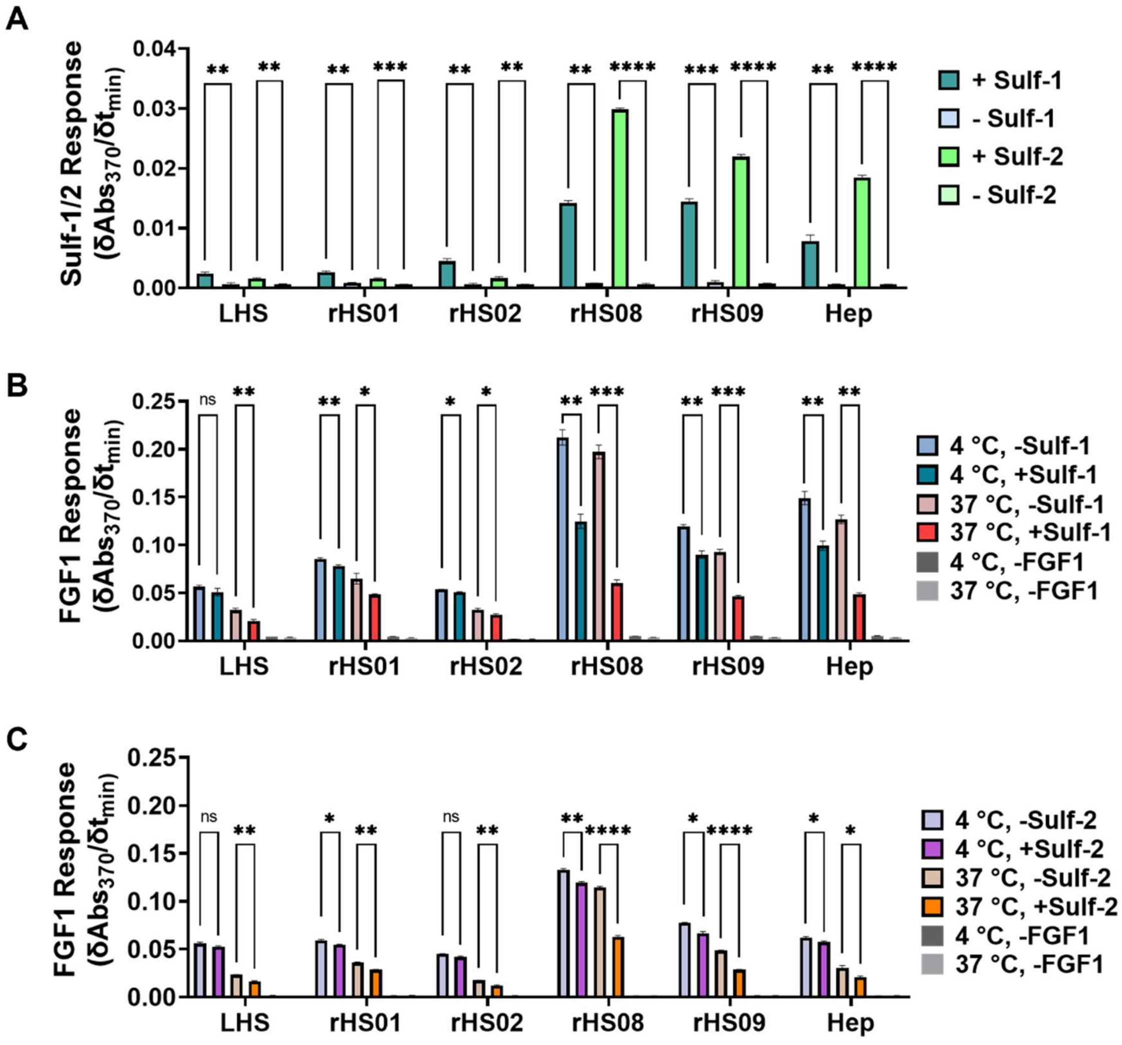

**Fig. S6. Sulf-1 and Sulf-2 binding to PG-mimetics and their effect of their activity on FGF1 association.** Raw data used for Figure 5A, 5C, and 5D. All samples used 100 uL volume, 100 ng/well $rHS_x$-BSA, 0.05 ug/uL Sulf-1/2 xCM. (*A*) 1:200 α-Sulf-1, 1:500 α-Rb HRP-IgG; 1:200 α-Sulf-2, 1:500 α-Ms HRP-IgG. (*B*) 25nM FGF1, 1:500 α-FGF1, 1:1000 α-Rb HRP-IgG. (*C*) 25nM FGF1, 1:500 α-FGF1, 1:1000 α-Rb HRP-IgG. (Bar graphs and error bars represent mean with SD, n = 3 independent experiments, *p*-values were determined using an unpaired Welch's t-test, $^{**}p < 0.01$, $^{***}p < 0.001$, $^{****}p < 0.0001$).

| Chemical Reagents | Source | Catalog No. |
|---|---|---|
| 1,8-diazabicyclo [5.4.0]undec-7-ene (DBU) | Sigma Aldrich | 139009-25G |
| 2-aminoacridone | Sigma Aldrich | 6627 |
| 3-azido-7-hydroxycoumarin | Carbosynth | FA31762 |
| 4-(Acetylamino)phenyl]imidodisulfuryl difluoride (AISF) | Sigma Aldrich | 901243 |
| Biotin-dPEG$_{11}$-Azide | Quanta Biodesign | 10784 |
| Carbazole | Ultra Scientific | HAH-022 |
| NHS-BCN | Sigma Aldrich | 744867-10MG |
| TAMRA-NHS | Invitrogen | C300 |
| Triton X-100 | Alfa Aesar | A16046 |
| Tween-20 | VWR | M147-4L |
| TMB substrate solution | VWR | 97063-666 |
| Phosphate Buffered Saline with Ca/Mg | Corning | 21-030-CM |
| **Biological Reagents** | **Source** | **Catalog No.** |
| Bovine Serum Albumin (BSA) | Spectrum | A3611 |
| Vascular Endothelial Growth Factor a (VEGF) | Thermo | PHC9394 |
| Fibroblast growth factor 1 | Abcam | Ab91374 |
| Fibroblast growth factor 2 | Peprotech | 100-18B |
| Fibroblast growth factor receptor 2α (IIIc) | R&D Systems | 712-Fr |
| BMP4 | Peprotech | 120-05 |
| Anti-VEGF | Invitrogen | P802 |
| Anti-FGF1 | Invitrogen | PA5-79249 |
| Anti-FGF2 | Millipore | 05-118 |
| Anti-BMP4 | Peprotech | 500-M121-500UG |
| Anti-mouse IgG HRP | Cell Signaling | 7076s |
| Anti-rabbit IgG HRP | Cell Signaling | 7074S |
| Heparin | Iduron | HEP001 |
| Heparinase I, II, III recombinantly purified | Gift from Jeffrey Esko lab | |
| Recombinant HS (01, 02, 08, 09, 29) | TEGA Therapeutics | rHS-01,02,08,09,29 |
| Streptavidin-HRP | Raybiotech | EL-HRP |

| | | |
|---|---|---|
| **Consumables** | **Source** | **Catalog No.** |
| Microtiter Plate Black clear-bottom high-binding 96-well | Greiner | 655097 |
| 3 kDa Centrifugal Spin Filters | Amicon | UFC500396 |
| 50 kDa Centrifugal Spin Filters | Amicon | UFC505096 |
| 25 kDa Dialysis Tubing | Spectra | 132126 |
| 1.5 mL Microcentrifuge Tube | Thermo Fisher | 5408129 |
| Microtiter Plate Black clear-bottom 96-well | Corning | 3915 |
| Microtiter Plate Clear 96-well | Corning | 3370 |
| Microtiter Plate EIA/RIA half-area high-binding 96-well | Corning | 3690 |
| PCR tube | Thermo Fisher | AB0337 |
| PD-10 Desalting Column | GE Life Sciences | 17085101 |

**Table S1.** Reagents and Consumables
